# Chemical Microenvironment of the Coral Gastrovascular Cavity and Its Associated Bacterial Diversity

**DOI:** 10.64898/2025.12.01.690986

**Authors:** Qingfeng Zhang, Elena Bollati, Michael Kühl

## Abstract

Coral gastrovascular cavities (GVCs) host diverse microorganisms and, as semi-closed compartments, are characterized by elevated nutrient concentrations and pronounced diel fluctuations in oxygen (O2) and pH. However, their broader chemical microenvironment and associated microbial activities remain poorly understood. Here, we provide direct evidence for active anaerobic metabolism in the GVC by measuring hydrogen (H2), nitric oxide (NO), and nitrous oxide (N2O) production, indicative of microbial fermentation, nitrogen fixation and denitrification. Changes in GVC H2 concentrations after feeding suggest that feeding modulates substrate availability, which in turn influences microbial activity in this compartment. In bleached corals, the GVC remained consistently hypoxic and acidic, and anaerobic metabolic processes persisted across the diel cycle. These patterns suggest a shift in energy and nutrient cycling pathways within the bleached coral holobiont. Together, our findings identify the GVC as a dynamic site of microbial activity that may play an important role in holobiont carbon and nitrogen cycling, with potential implications for nutrient balance and coral resilience.

## Introduction

Tropical, reef-building corals represent a multispecies assemblage of calcifying cnidarian polyp animals, their microalgal endosymbionts (in the dinoflagellate family Symbiodiniaceae), as well as multiple microbiomes associated with the animal tissue surface, the animal gastrovascular cavity (GVC), and the underlying coral skeleton forming the coral colony scaffold (Hughes et al., 2022; Voolstra et al., 2024). While the microenvironmental conditions in these compartments of the coral holobiont can be very different (e.g. Kühl et al., 1995; Kühl et al., 2008; Agostini et al., 2012; Lichtenberg et al. 2016; Kühl et al., 2024), most ecophysiological studies of corals are done at the level of whole coral colony fragments. Likewise, there is increasing evidence for differential composition of microbiomes in different coral holobiont compartments (e.g. Sweet et al., 2011; Apprill et al., 2016; Marcelino et al., 2018; Ricci et al., 2023), and there is a rising awareness among coral biologists of the need for higher spatial resolution studies to unravel functional links between the coral host and different microbial communities in the holobiont (van Oppen and Raina, 2023).

The Symbiodiniaceae within coral tissues not only shape the O2 and pH microenvironment of corals via the interplay of symbiont photosynthesis, host and symbiont respiration (Kühl et al., 1995), but also fundamentally determine the coral holobiont’s energy acquisition (Ianniello et al., 2025). Upon coral bleaching, i.e., the breakdown of this mutualistic symbiosis due to environmental stressors, the loss of algal symbionts disrupts the primary energy supply of the host and forces corals to shift their energy acquisition toward heterotrophy (Grottoli et al., 2006; Houlbrèque and Ferrier-Pagès, 2009; Meunier et al., 2019). Such changes may also profoundly alter the nutrient composition and redox state within the GVC. Consequently, its microbial community composition and activity may also be altered, potentially affecting holobiont recovery and survival. Yet, how different compartments and associated microbes respond to bleaching remains poorly understood, leaving a critical gap in our knowledge of coral holobiont function under stress.

The GVC of corals plays a central role for their biology, e.g., as a site for food digestion, nutrient uptake and sexual reproduction, as a pathway for departing or arriving symbionts during coral bleaching and recovery, as well as for resource sharing between neighboring polyps (Hughes et al., 2022; Kühl et al., 2024). The semi-enclosed nature, small volume and large internal tissue surface area of the GVC leads to strong spatio-temporal variations in O2 concentration and pH depth gradients in the GVC between light and dark conditions affecting e.g. the internal carbonate chemistry and calcification process in corals (Agostini et al., 2012; Jokic et al., 2012; Cai et al., 2016; Bove et al., 2020; Dellisanti et al., 2024; Crovetto et al., 2024).

While the chemical conditions in the GVC of some corals switch from more uniform high pH and hyperoxia in light to low pH and hypoxia/anoxia in darkness (Zhang et al., in review), other species with deeper GVCs such as *Galaxea fascicularis*, *Dipsastraea favus* and *Lobophyllia hemprichii* can exhibit strong stratification of their chemical microenvironment rendering deeper parts of the GVC hypoxic or anoxic even in light (Agostini et al., 2012; Bollati et al., 2024). Such spatiotemporal dynamics potentially allow for the presence of anaerobic or microaerophilic microbes in the coral GVC, however, so far direct evidence of anaerobic processes in the GVC via microsensor measurements of e.g. H2 (indicative of fermentation or N2 fixation), NO or N2O (indicative of denitrification), or H2S (indicative of sulfate reduction) is lacking.

In this study, we used O2, pH, N2O, NO and H2S microsensors to assess the chemical microenvironment and its dynamics in the GVC of healthy and bleached corals. We combined such measurements with microvolume sampling and subsequent assessment of microbial diversity in the GVC of individual coral polyps using amplicon sequencing. Our study represents the hitherto most detailed characterisation of the GVC microenvironment and its implications for microbial diversity and function in this central compartment of the coral holobiont.

## Materials and methods

### Coral husbandry

Colonies of *Caulastrea curvata* were obtained from DeJong Marine Life (Netherlands) and maintained in a recirculating aquarium system at the Marine Biology Section, University of Copenhagen (Helsingør, Denmark). Upon delivery, three polyps appeared visibly bleached (possibly due to low-temperature stress during transport). Healthy and bleached corals were kept under controlled conditions: temperature 25°C, salinity 35, and photon irradiance (400–700 nm) of 150 μmol photons m^−2^ s^−1^, with a 12 h light:12 h dark cycle for at least 7 days before measurement. The same sets of healthy *C. curvata* polyps (1–3) and bleached *C. curvata* polyps (1–3) were used for O2, pH, H2, NO, and GVC microsample analyses. For N2O measurements, a separate set of healthy *C. curvata* polyps from a different colony was used. Two fragments of *Goniastrea* sp. were collected from reef flat of Heron Island (Great Barrier Reef, Australia; permit n. G24/49877.1) in May 2024 using a hammer and chisel, and kept at an outdoor flow-through aquarium flushed with seawater from the reef for 7 days before measurement.

### Microsensor measurements

Microsensors for O2, pH, H2, H2S, NO and N2O (Unisense A/S, Denmark) with a tip diameter of 50 μm were used to measure chemical dynamics in the coral GVC microenvironment.The microsensors (and a reference electrode for pH measurements; Ag/AgCl, Unisense A/S, Denmark) were connected to a multichannel microsensor meter (fx-6 UniAmp, Unisense A/S, Denmark) and mounted on a motorized micromanipulator (Unisense A/S, Denmark) that enabled precise vertical movement at µm-scale resolution. The microsensor meter and micromanipulator were connected to a PC with dedicated software for data acquisition and sensor positioning (SensorSuite Profiler v3.4, Unisense A/S, Denmark). Each microsensor was calibrated at experimental temperature and salinity prior to measurement.

The O2 microsensors were linearly calibrated from sensor signal readings (in pA) using a two-point method in 100% air-saturated seawater and anoxic water (using a sodium ascorbate solution). The pH microsensors were calibrated from sensor readings (in mV relative to the reference electrode) in standard buffer solutions of pH 4.0, 7.0 and 10.0 (Hach, USA). The H2 microsensors were linearly calibrated from sensor readings (in pA) in seawater from the experimental aquarium (0 μM H2) and seawater flushed with a gas mixture of 5% H2 and 95% N2 gas, yielding a H₂ concentration of 33.4 μM at experimental temperature and salinity according to tabulated values of gas solubility in water (unisense. com). The NO microsensor was calibrated using a solution containing 0.1 M H2SO4 and 0.1 M KI, which was flushed with N2 gas to remove dissolved O2. 200 μL of 1 mM NaNO2 solution was added stepwise into the 200 mL calibration solution. Each addition generated 1 μM NO, and the NO microsensors were linearly calibrated from microsensor readings (in pA) in 0, 1, 2 and 3 µM NO solutions. The N2O microsensor was linearly calibrated from readings (in pA) in seawater from the experimental aquarium (0 μM N2O) and 100 μM N2O solution prepared by adding 0.95 mL of N2O-saturated solution to 199.05 mL seawater. The H₂S microsensor was linearly calibrated from readings (in pA) using a pH 4 buffer solution (salinity 35) flushed with N₂ gas to remove oxygen. The 0 µM solution was the buffer, a 100 µM H₂S solution was prepared by adding 0.1 mL of 0.01 M Na₂S to 9.9 mL buffer; and a 50 µM H₂S solution was obtained by mixing equal volumes of the 0 and 100 µM solutions.

The positioning of the microsensor tip relative to the coral mouth was monitored using a digital microscope (Dinolite, AnMo Electronics, Taiwan). The sensor tip was manually aligned to the center of the coral mouth, and was then lowered in 20–50 μm steps using the motorized micromanipulator until coral tissue contraction was observed, which was defined as the bottom of the GVC. Measurements were initiated after the sensor signal had stabilized for 5 min.

During microsensor measurements, the coral fragment was placed in a custom-designed flow chamber with a constant laminar flow (1 cm s ^-1^) of thermostated and oxygenated seawater (kept at 25°C and a salinity of 35) supplied from a supporting aquarium, as previously described by Brodersen et al. (2014) and Zhang et al. (2025). During light measurements, the coral fragment was illuminated by a white LED lamp (KL 2500 LED, Schott, Germany) providing a photon irradiance (400-700 nm) of 100 μmol photons m^-2^ s^-1^ (low light) or 200 μmol photons m^-2^ s^-1^ (high light). Photon irradiance was quantified with a photon irradiance meter (ULM-500, Heinz Walz, Germany) equipped with a spherical micro quantum sensor (US-SQS/L, Heinz Walz, Germany). For H2 and NO measurements on healthy polyps, high light measurements were only conducted if H₂ or NO was detected under low light conditions.

Prior to the onset of experiments, coral fragments were fed with coral food (ReefPearls 5–200 µm, DVH Aquatic, Netherlands) 20–24 h prior to measurements. For H2 monitoring after feeding, *C. curvata* was kept under a 12 h:12 h light–dark cycle for 36–48 h, with measurements conducted at three time points. For continuous H₂ measurements in the GVC of *Goniastrea* sp. (see setup details below), coral food was delivered directly to the oral disks of target polyps using a pipette, and the microsensor tip was positioned at approximately 50% of the cavity depth.

### Microvolume sample collection, DNA extraction and 16S rDNA metabarcoding

For amplicon sequencing-based microbiome analysis, GVC fluid samples were collected after microsensor measurements. During sampling, coral fragments were placed in a glass container that had been rinsed with 70% ethanol and filled with 0.22 μm-filtered seawater to avoid cross-contamination. Microvolume samples of GVC fluid were collected from *C. curvata* polyps using a sterile 1 mL syringe equipped with a low dead-space microneedle (34G, TSK, Canada) as described in Bollati et al. (2024). Seawater samples (100 µL) were also collected from the sampling container, aquarium tank, and flow chamber using the same method. All samples were transferred into UV-crosslinked (1 h) 1.5 mL sterile microcentrifuge tubes, and stored at −70 °C until processing.

DNA extraction was performed in a UV-sterilized hood using a low-input (10 µL) physical lysis protocol, following Bramucci et al. (2021) and Bollati et al. (2024). The V3–V4 hypervariable region of the 16S rRNA gene was amplified using Illumina fusion primers (341 F: TCGTCGGCAGCGTCAGATGTGTATAAGAGACAG CCTAYGGGRBGCASCAG and 805 R: GTCTCGTGGGCTCGGAGATGTGTATAAGAGACAG GGACTACNNGGGTATCTAAT).

Each 25 µL PCR reaction contained 5 µL template DNA, 12.5 µL KAPA HiFi HotStart ReadyMix (Roche, Switzerland), 0.5 µL of each 10 µM primer, and 6.5 µL PCR water. PCR amplification was performed with an initial denaturation at 98 °C for 2 min, followed by 35 cycles of 98 °C for 30 s, 55 °C for 30 s, and 72 °C for 30 s, with a final elongation at 72 °C for 10 min. Amplicons were visualized by agarose gel electrophoresis and sent to the Australian Genome Research Facility (Melbourne, Australia) for library preparation and sequencing on the Illumina NextSeq platform (300 bp paired-end). Two extraction negatives were included and sequenced alongside the samples.

### Sequencing data processing

Demultiplexed sequences were trimmed to remove adapter and primer sequences using cutadapt (v4.4) (Martin, 2011), and subsequently processed in R (v4.1.1) with the DADA2 pipeline (v1.22) (Callahan et al., 2016). Forward and reverse reads were truncated at 250 bp, and the maximum number of expected errors was set to 2. Filtered reads were denoised, merged, and screened for chimeras to infer amplicon sequence variants (ASVs). Taxonomic classification of ASVs was performed against the SILVA reference database (v138.1) (Quast et al., 2013). ASVs in extraction blanks and ASVs identified as eukaryotic, chloroplast, or mitochondrial sequences were removed. Rarefaction curves were generated using the vegan package (v2.6-4) (Oksanen et al., 2022) to verify sufficient sequencing depth, and no rarefaction was applied.

Functional prediction of microbial communities was performed using PICRUSt2 (v2.5.0) (Douglas et al., 2020) to generate KEGG Ortholog (KO) functional profiles. Subsequent analyses focused on anaerobic metabolic pathways, including denitrification, nitrogen fixation, hydrogen production and consumption, sulfate reduction and sulfide oxidation pathways (Table S1). ASVs corresponding to target KOs were identified, and the predicted abundance of each function was calculated as the relative abundance of ASVs assigned to any of the corresponding KOs in each sample. The predicted abundance of the antioxidant enzyme catalase was compared between samples from healthy and bleached corals; while predicted denitrification and hydrogen metabolism functions were compared among groups defined by different NO and H₂ concentrations measured by microsensors.

Alpha diversity was calculated from the ASV table in *phyloseq* v1.52 as Shannon’s H index and visualized as its exponential (eH). Differences in alpha diversity among SampleType groups were compared using the Kruskal–Wallis test, followed by pairwise Wilcoxon rank-sum tests with Benjamini–Hochberg correction. Beta diversity was assessed using Bray–Curtis dissimilarity, and microbial community composition was visualized via principal coordinates analysis (PCoA) using *phyloseq*. Differences in community composition were tested via PERMANOVA (adonis2, 999 permutations), and homogeneity of dispersion was confirmed using *betadisper* with ANOVA in the R package vegan v.2.7-1 (Oksanen et al., 2025). Differential abundance was evaluated pairwise using ALDEx2 v1.26.0 (Fernandes et al., 2013) for taxa aggregated at the family level to identify families exhibiting significant differences in relative abundance across sample groups. Taxa were considered significant if they had an unadjusted p-value < 0.05 and the 95% confidence interval of effect size overlap was less than 20%. Relative abundances of selected predicted functional genes were compared between groups using the Wilcoxon rank-sum test. All analyses were performed in R v4.4.3.

## Results

### Chemical dynamics in the coral GVC

In the GVC of healthy corals, O2 concentrations reached 350–400 μmol O2 L^-1^ under high light, 250–300 μmol O2 L^-1^ under low light, and dropped to below 100 μmol O2 L^-1^ in the dark (Figure 1A). Spatial profiling revealed that under light conditions, O2 levels were higher in the upper region of the GVC and then decreased with depth, while O2 concentration declined from from the mouth opening into the GVC during darkness.

**Figure 1.**
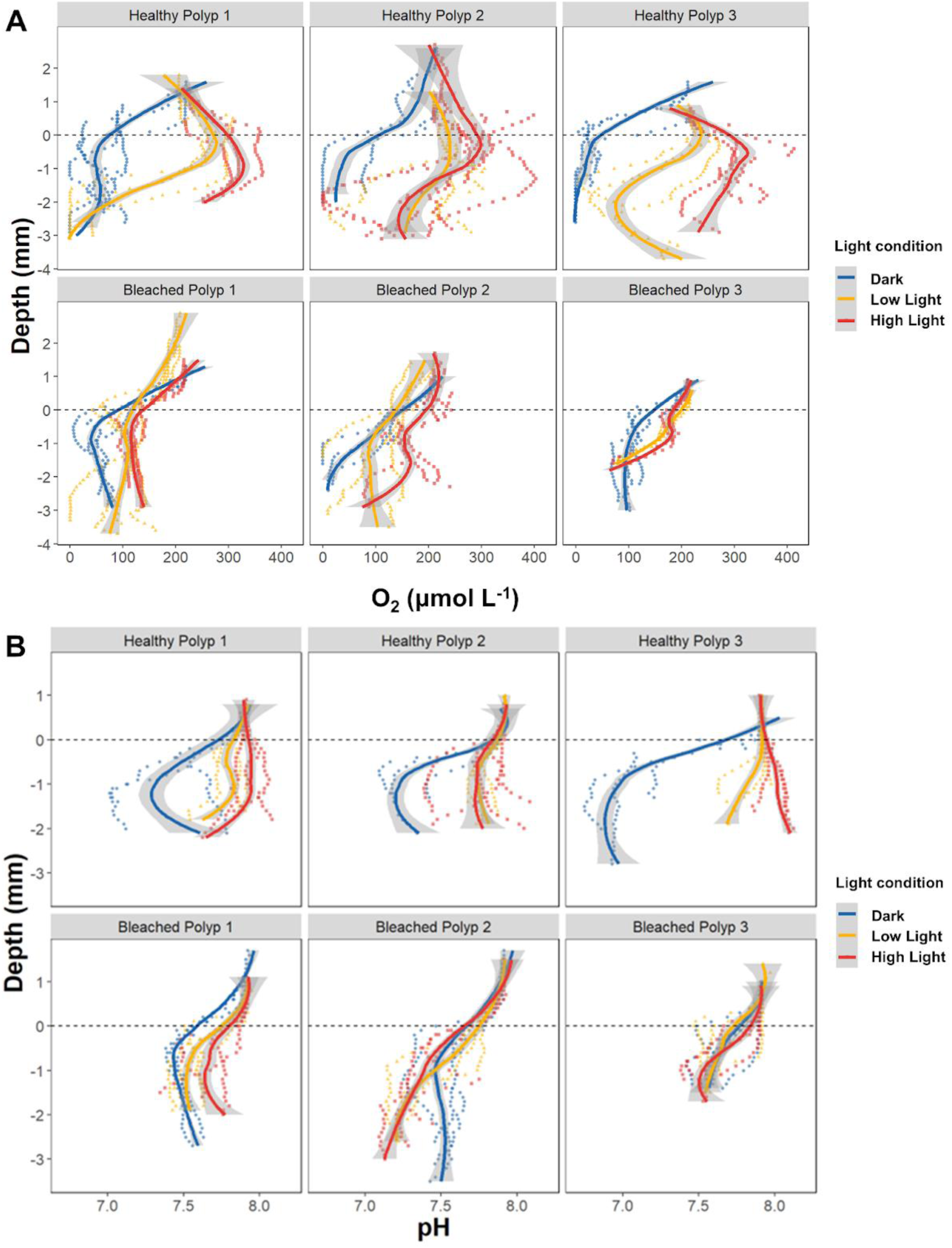
Depth profiles of O2 concentration (A) and pH (B) in the gastrovascular cavity of healthy and bleached *C. curvata* polyps in darkness and under an incident photon irradiance (400-700 nm) of 100 µmol photons m^-2^ s^-1^ and 200 µmol photons m^-2^ s^-1^. The dashed lines indicate replicate measurements (n = 3), while the continuous lines indicate LOESS-smoothed trends with 95% confidence intervals (grey shading). Zero depth represents the level of the coral mouth opening, while negative depth values indicate positions inside the GVC.

However, technical replicate measurements on the same coral polyp also showed pronounced fluctuations under constant light conditions. For example, in Healthy Polyp 2, the O2 concentration shortly dropped to anoxic level under high light. In comparison, bleached corals showed consistently lower O_2_ levels (below 200 μmol O_2_ L^-1^), with only slight increases under light relative to dark conditions.

Similar to O₂ dynamics, pH variations were mainly governed by the light-dark cycle. In the healthy corals, pH in the GVC varied by > 1 pH unit, i. e., from < pH 7.0 in the dark to > pH 8.0 under high light. In bleached corals, pH remained consistently lower than ambient seawater pH under both dark and light conditions (Figure 1B).

Despite low pH and hypoxic/anoxic conditions in the GVC of dark incubated, healthy or bleached *C. furcata* corals, we did not detect any H₂S, which would be indicative of active sulfate reduction in the GVC. Instead, we documented pronounced H₂ dynamics in the GVC of dark and light incubated corals (Figure 2, Figure 3). In Healthy Polyp 1 and 2, the maximum H_2_ concentration was 0.05-0.08 μmol H_2_ L^-1^ in the dark and decreased to 0 under low light conditions. Healthy Polyp 3 showed higher H2 production, with a maximum H_2_ concentration of 0.77 μmol H_2_ L^-1^ in the dark, remaining relatively high (0.51 μmol H_2_ L^-1^) under low light, and then decreasing to below 0.04 μmol H_2_ L^-1^ under high light. In bleached corals, the overall H2 concentrations were higher than those in healthy polyps, the maximum H2 concentration in the dark ranged from 0.30 to 0.76 μmol H_2_ L^-1^. Furthermore, sustained H_2_ production in the GVC of bleached corals was observed under illumination, even under high light, with concentrations reaching 0.11 to 0.22 μmol H L^-1^.

**Figure 2.**
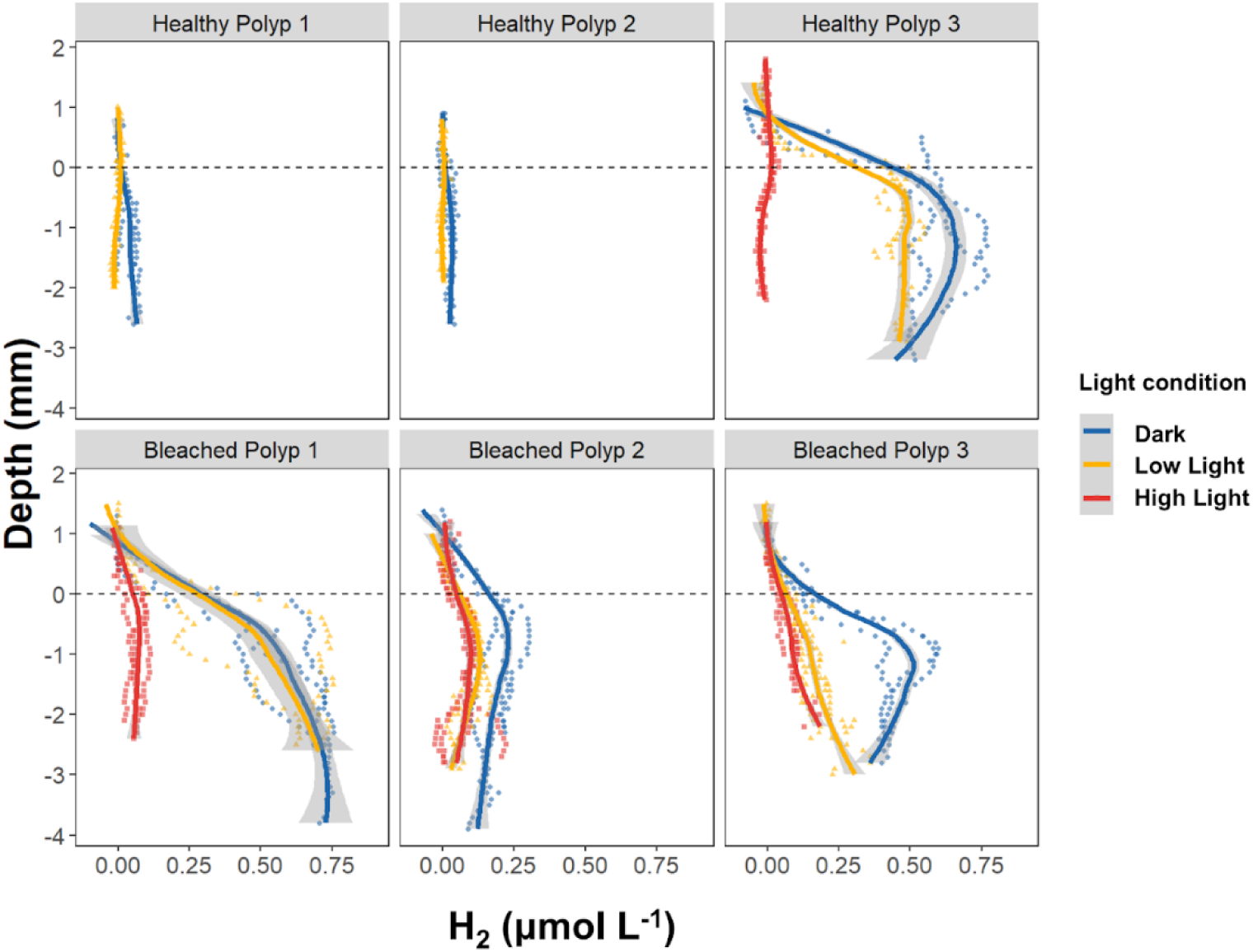
Depth profiles of H2 concentration in the gastrovascular cavity of healthy and bleached *C. curvata* polyps in darkness and under an incident photon irradiance (400-700 nm) of 100 µmol photons m^-2^ s^-1^ and 200 µmol photons m^-2^ s^-1^. The dashed lines indicate replicate measurements (n = 3), while the continuous lines indicate LOESS-smoothed trends with 95% confidence intervals (grey shading). Zero depth represents the level of the coral mouth opening, while negative depth values indicate positions inside the GVC.

**Figure 3.**
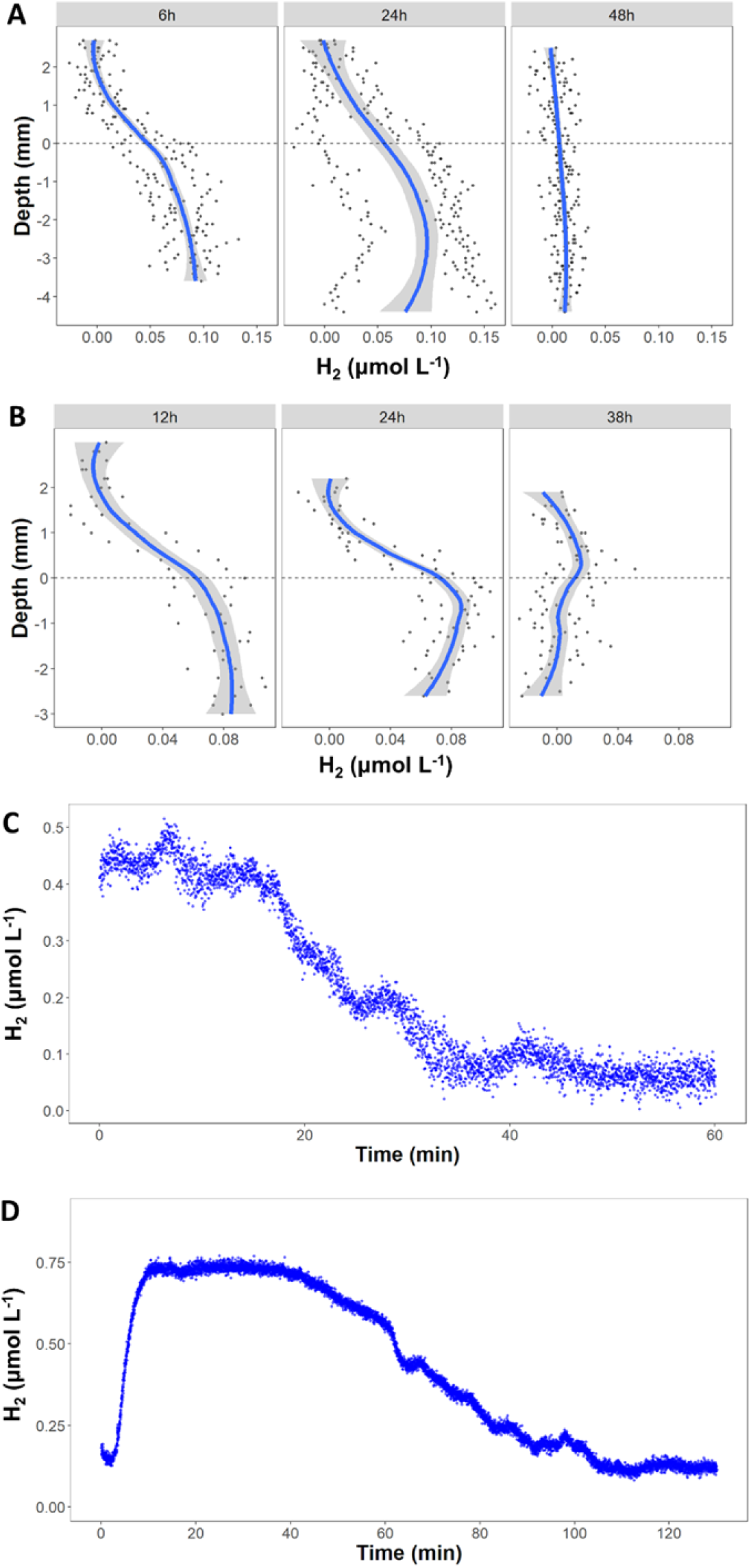
Temporal dynamics of H2 concentration in the gastrovascular cavity after feeding under dark conditions. (A-B) H2 depth profile in the healthy *C. curvata* polyps. (C-D) Continuous measurement in *Goniastrea* sp.; the microsensor tip was positioned at 50% of polyp depth. (C) measurement started after H2 concentration stabilized; (D) measurement started after light off.

We also monitored the H2 concentration dynamics in the GVC after feeding. In *C. furcata* polyps, the H_2_ concentration remained elevated (0.10 μmol H_2_ L^-1^ and 0.08 μmol H_2_ L^-1^ in replicate 1 and 2, respectively) for at least 24 h, before returning to zero at 36–48 h (Figure 3A, B). In *Goniastrea* sp. polyps, the continuous measurements revealed rapid changes in H_2_ concentration following feeding (Figure 3C, D). H_2_ level increased to 0.5 μmol H_2_ L^-1^ in replicate 1 (Figure 3C); in replicate 2, the H_2_ concentration increased to 0.75 μmol H_2_ L^- 1^ within 10 minutes after light off and began to decrease after 1 h (Figure 3D).

The intermediate products of the denitrification pathway, NO and N2O, were also detected under hypoxic conditions in the GVC. Some samples exhibited very high NO concentrations, reaching up to 1.2 μmol NO L^-1^ in Healthy Polyp 2 and 0.95 μmol NO L^-1^ in Bleached Polyp 3 (Figure 4). In healthy corals, NO followed the same light-dependent pattern as H2, decreasing with increasing irradiance: In Healthy Polyp 1, the maximum NO concentration decreased from 0.22 μmol NO L^-1^ in the dark to undetectable levels under low light, and in Healthy Polyp 2, it decreased from 1.2 μmol NO L^-1^ in the dark to 0.44 μmol NO L^-1^ under high light. Compared with the other polyps, Healthy Polyp 3 and Bleached Polyp 1 exhibited consistently low NO concentrations, remaining below 0.06 μmol NO L^-1^ across all light conditions. We observed high variability in NO dynamics among technical replicates. For example, under high light condition: in Bleached Polyp 2, two technical replicates exceeded 0.3 μmol NO L^-1^, whereas one remained below 0.05 μmol NO L^-1^; in Bleached Polyp 3, two replicates were lower than 0.06 μmol NO L^-1^, while one reached as high as 0.95 μmol NO L^-1^. In contrast, the N2O concentrations remained relatively stable in the GVC, ranging from 0.08 to 0.20 μmol N2O L^-1^ in healthy polyps in the dark (Figure 5).

**Figure 4.**
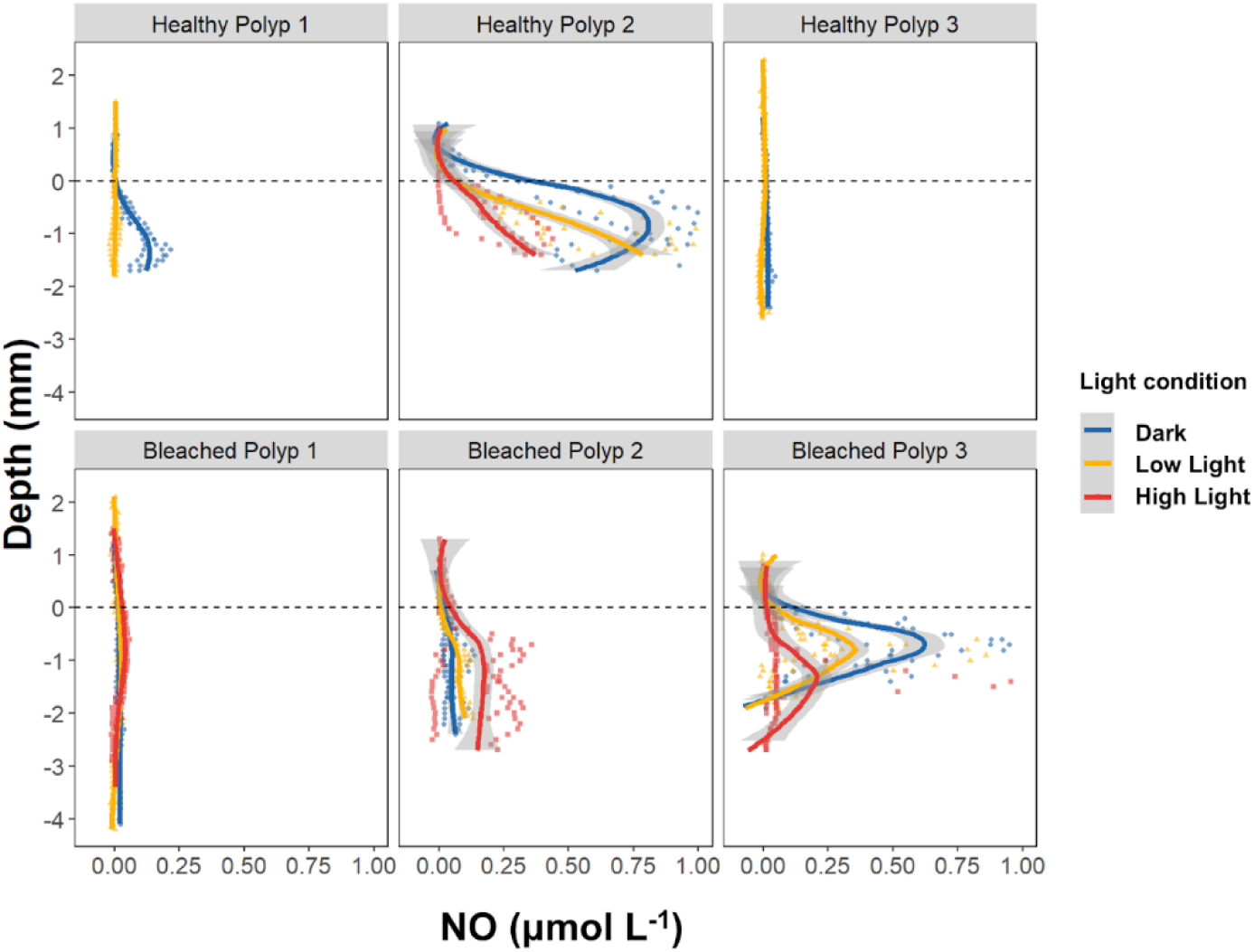
Depth profiles of NO in the gastrovascular cavity of healthy and bleached *C. curvata* polyps in darkness and under an incident photon irradiance (400-700 nm) of 100 µmol photons m^-2^ s^-1^ and 200 µmol photons m^-2^ s^-1^. The dash lines indicate replicate measurements (n = 3), while the continuous lines indicate LOESS-smoothed trends with 95% confidence intervals (grey shading). Zero depth represents the level of the coral mouth opening, while negative depth values indicate positions inside the GVC.

**Figure 5.**
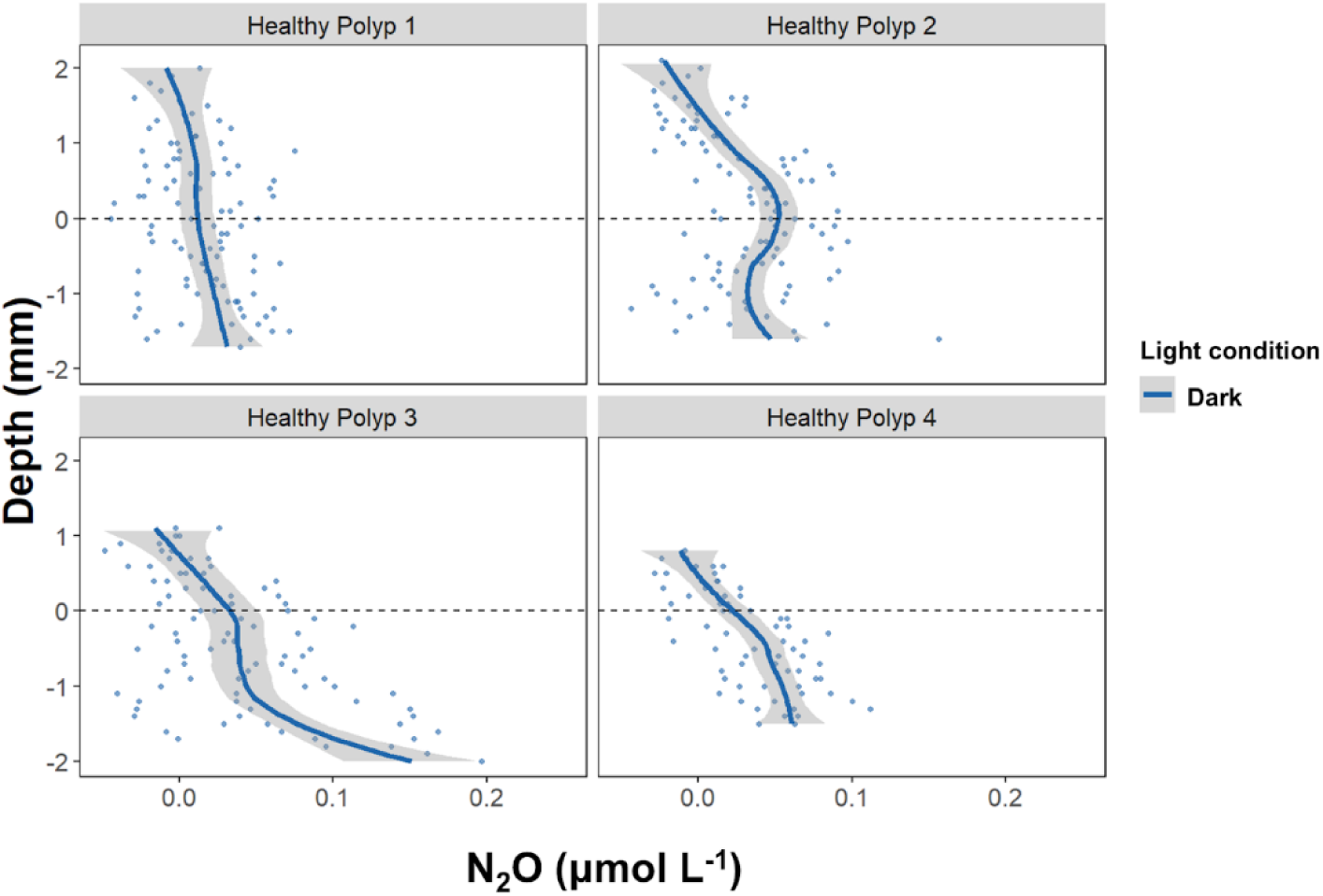
Depth profiles of N2O in the gastrovascular cavity of healthy *C. curvata* polyps under dark conditions. The dash lines indicate replicate measurements (n = 3), while the continuous lines indicate LOESS-smoothed trends with 95% confidence intervals (grey shading). Zero depth represents the level of the coral mouth opening, while negative depth values indicate positions inside the GVC.

### Microbial diversity in the coral GVC

Microvolume sampling and subsequent DNA extraction and amplicon sequencing was successful in all sampled polyps, yielding (1.2–6.3) ×10^5^ decontaminated reads in GVC samples, aquarium samples and flow chamber samples, and (2.1–3.5) ×10^4^ reads in filtered seawater samples. PCoA based on Bray–Curtis dissimilarities revealed clustering patterns among five sampling groups (Fig. 6A), which was further supported by relative abundance profiles of bacterial families (Fig. S2). GVC-associated samples (healthy and bleached polyps) generally clustered separately from environmental samples, and healthy and bleached polyps were mostly distinct. However, one healthy polyp sample (Healthy Polyp1) was positioned closer to the bleached cluster, suggesting some overlap in community composition. PERMANOVA confirmed that microbial community composition differed significantly between groups (*adonis2*: F = 1.25, R^2^ = 0.384, p = 0.046). Multivariate dispersion analysis indicated no significant differences in within-group variability (*betadisper*: F = 0.85, p = 0.594), confirming that the observed PERMANOVA result was not driven by heterogeneous dispersion.

**Figure 6.**
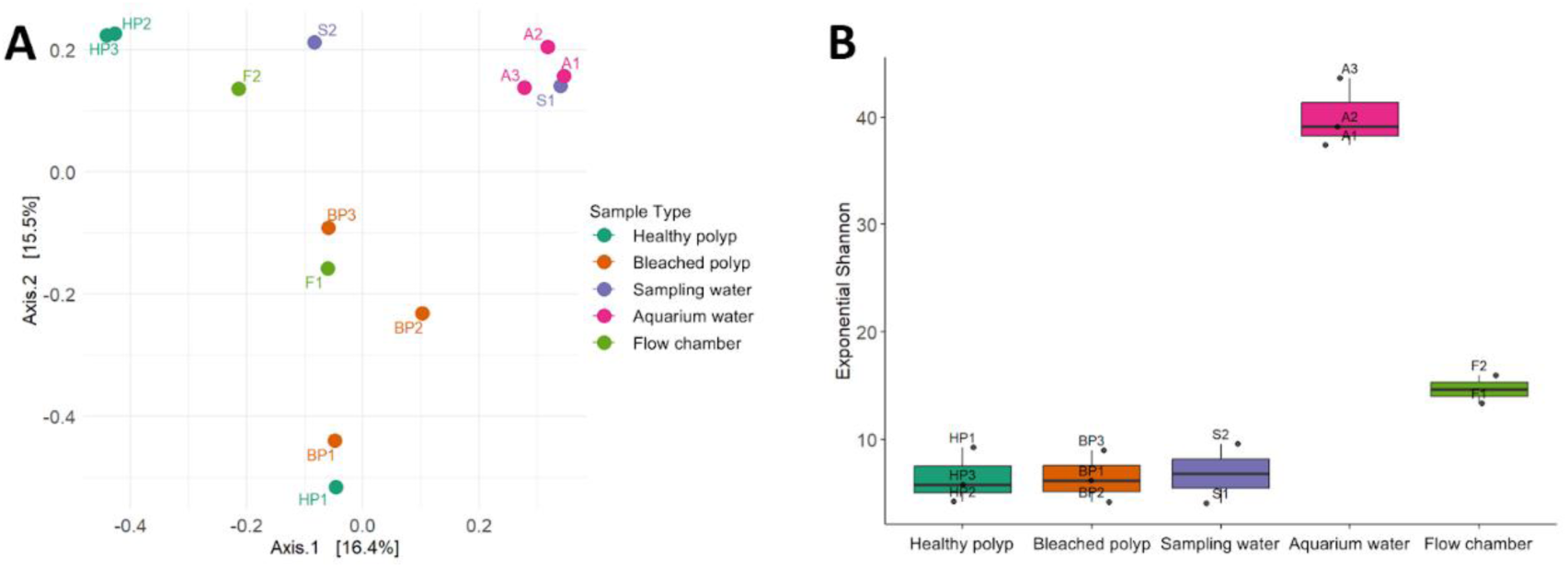
Microbial diversity in the gastrovascular fluid of healthy and bleached *C. curvata* polyps and in environmental samples. (A) Principal coordinates analysis (PCoA) based on Bray–Curtis dissimilarities showing differences in bacterial community composition among sample types. (B) Alpha diversity represented as the exponential of the Shannon index (e^H^).

Alpha diversity, assessed as Shannon’s H (exponential), was broadly comparable among healthy polyps, bleached polyps, sampling water, and flow chamber samples, while aquarium water samples exhibited higher diversity (Figure 6B). However, Kruskal–Wallis testing did not detect significant differences across groups (χ^2^ = 9.08, p = 0.059), and pairwise Wilcoxon rank-sum tests with Benjamini–Hochberg correction did not reveal significant contrasts (all adjusted p > 0.33). These results indicate that overall alpha diversity is largely comparable across different sampling groups, reflecting similar microbial richness and evenness.

To identify the taxa contributing to the observed differences in community composition, we examined the relative abundances of bacterial families between healthy and bleached coral polyps (Figure 7A). Bleached polyps showed a higher relative abundance of Staphylococcaceae, whereas healthy polyps were enriched in Micrococcaceae. Given the variation in NO and H2 concentrations within the GVC among individual coral polyps, we further tested whether these microenvironmental gradients corresponded to shifts in family-level bacterial composition (Figure 7B, C). Polyps with low H2 concentrations exhibited higher abundances of Rhizobiaceae and Rhodobacteraceae. Polyps with low NO concentrations were enriched in Pseudoalteromonadaceae and Spongiibacteraceae, whereas Xanthobacteraceae and Alcanivoracaceae were more abundant in high-NO polyps.

**Figure 7.**
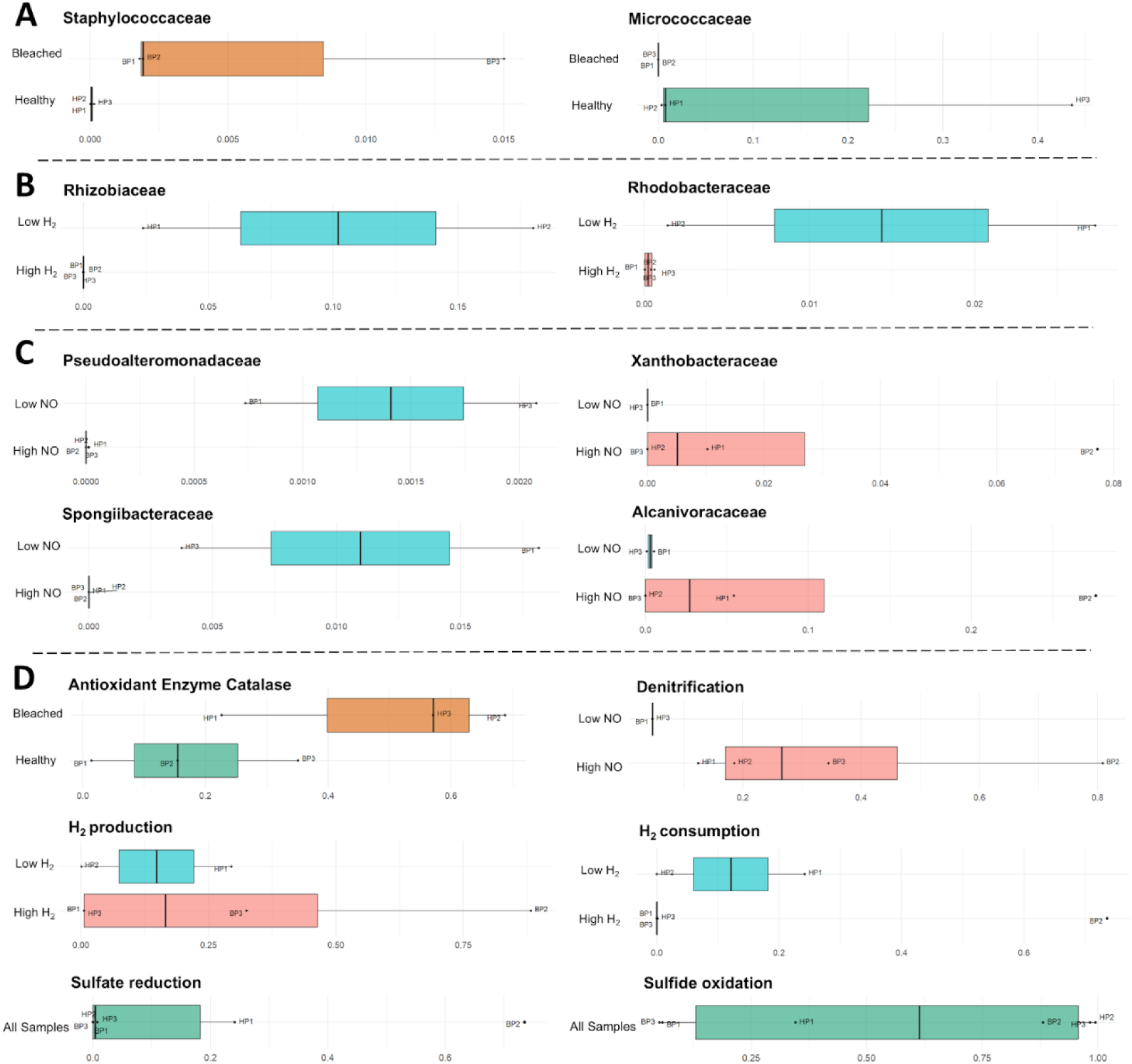
Relative abundance of bacterial families and predicted functions across different sample groups. (A–C) Family-level relative abundance of bacterial taxa showing significant differences between groups: (A) Healthy or bleached coral polyps; (B) High or low GVC H₂ concentration; (C) High or low GVC NO concentration. (D) Relative abundance of predicted microbial functions based on KEGG Orthologs corresponding to target pathways.

We next evaluated the functional profiles of the GVC-associated microbiomes using predicted gene abundances, focusing on antioxidant activity, denitrification, hydrogen and sulfur metabolism (Figure 7D). Importantly, we note that these predictions are based on the taxonomic profiles present in each sample, and therefore do not represent the true presence or absence of metabolic pathways or genes of interest. The predicted abundance of the antioxidant enzyme catalase appeared to be higher in healthy polyps compared with bleached polyps (49.5 ±24.0% and 17.3 ± 16.9%, respectively), although this difference was not statistically significant (p = 0.20). For denitrification-related genes, polyps with low NO concentrations exhibited lower predicted abundances (<5%) than those with high NO concentrations (12.5–80.8%), but the difference did not reach statistical significance (p = 0.13). Predicted functions related to H2 production and consumption were comparable among groups, with no statistically significant differences detected.The predicted *nifH* gene (associated with nitrogen fixation) showed a relative abundance of 3.2% in Bleached Polyp 2, while its abundance was below 0.1% in all other samples. We found relatively high abundances of predicted sulfate-reducing genes in Healthy Polyp 1 and Bleached Polyp 2, accounting for 24.2% and 73.1% of the community, respectively, whereas all other polyps showed levels below 1%. Predicted sulfide-oxidizing genes were generally more prevalent, reaching 5–6% in Bleached Polyps 1 and 3, 34.6% in Healthy Polyp 1, and as high as 88–99% in the other polyps.

## Discussion

Our microsensor measurements reveal that the coral gastrovascular cavity (GVC) is a highly dynamic chemical microenvironment that is affected by different microbial processes. Combined with taxonomic analysis of the GVC-associated microbiome, our findings highlight the tight coupling of host physiology, symbiont photosynthesis, and microbial metabolism in shaping microenvironmental conditions within coral polyps.

The “high light” and “low light” levels (200 and 100 μmol photons m^-2^ s^-1^, respectively) were chosen relative to the aquarium environment (150 μmol photons m^-2^ s^-1^) and are far lower than the high light levels experienced in natural reef habitats. Nevertheless, despite the relatively low light levels, pronounced diel dynamics in O2 and pH were observed in healthy polyps. From light to dark, the GVC of healthy polyps shifted from hyperoxic to hypoxic, with a substantial drop in pH. In contrast, O2 and pH profiles in the GVC of bleached coral polyps consistently exhibited hypoxic and acidic conditions under dark and different light conditions, reflecting the loss of O2 production by Symbiodiniaceae. These altered chemical conditions between bleached and healthy polyps likely reshape the balance between aerobic and anaerobic metabolism within the holobiont. In bleached polyps, the production of H2 and NO persisted under illumination, indicating that anaerobic processes such as fermentation and denitrification are sustained across the diel cycle. This persistence suggests a shift in energy and nutrient cycling pathways in the bleached coral holobiont. Consistent with this, we observed a marked decline in aerobic bacteria (Micrococcaceae) in bleached polyps. In contrast, the high predicted abundance of the antioxidant enzyme catalase in two out of three healthy polyps suggests that microbes associated with healthy corals may possess greater potential for ROS detoxification in response to photosynthesis-associated oxidative stress.

Beyond the pronounced changes in the GVC chemical microenvironment between healthy and bleached polyps, the collapse of the coral-algae symbiosis also induces other changes in host physiology that significantly influence the associated microbiome. Microbial community restructuring during bleaching has been widely reported (Bourne et al., 2008; Littman et al., 2011; Pogoreutz et al., 2018; Pootakham et al., 2018; Gardner et al., 2019; Kusdianto et al., 2021; Sun et al., 2022). This restructuring likely arises from shifts in energy metabolism following symbiont loss, and from a reduction in host immune function that reduces the host’s ability to regulate its microbiome (Pinzón et al., 2015). Although different coral species vary in the stability or flexibility of their bacterial community structure (Pogoreutz et al., 2018; Ziegler et al., 2019), a common pattern across studies is a shift toward higher abundances of heterotrophic and potentially pathogenic bacteria in bleached corals (Littman et al., 2011; Sun et al., 2022). We observed a higher relative abundance of Staphylococcaceae in the GVC of bleached polyps. Members of this family include opportunistic pathogens associated with host stress (Jordan et al., 1980; Lowy, 1998; Dinges et al., 2000; Otto, 2009), which may reflect these physiological changes in the coral host. This shift suggests that the GVC becomes a niche more permissive to opportunistic pathogens, potentially increasing the susceptibility of bleached corals to subsequent disease (Muller and Baum 2018). Environmental stress can also directly shape microbial communities, for example through changes in temperature or elevated nutrients (Thurber et al., 2009; Ziegler et al., 2019). However, in this study, we did not induce bleaching under controlled experimental conditions, so we cannot disentangle the direct effects of environmental variation.

The detection of H2, NO and N2O provides direct evidence of active anaerobic microbial processes within the GVC. These pathways represent key components of carbon and nitrogen turnover in both environmental and host-associated microbial communities. Fermentative H2 production reflects microbial degradation of labile organic carbon and generates organic acids and alcohols that can fuel downstream anaerobic metabolisms (Gibson et al., 1993). In principle, hydrogen gas production could also indicate N2 fixation, where H2 can be a byproduct (Bergersen, 1966). Denitrification reduces nitrate and nitrite to gaseous nitrogen species (NO, N2O and N2), thereby removing bioavailable nitrogen from the holobiont and helping to maintain the C–N balance, which is crucial for the stability of the coral–algae symbiosis (Rädecker et al., 2015; Tilstra et al., 2019; El-Khaled et al., 2021; Glaze et al., 2022). These anaerobic pathways are shaped not only by oxygen availability, but also by the local concentrations of organic carbon and inorganic nitrogen substrates (Chubukov et al., 2014). Because the GVC is the primary site of food acquisition and digestion, feeding directly enriches the cavity with labile organic matter, providing substrates that support fermentation and denitrification. Previous studies of the coral GVC identified bacterial taxa with known sulfate reduction and sulfide oxidation potential (Agostini et al. 2011, Bollati et al. 2024), a finding in line with our functional predictions of sulfate-reduction and sulfide-oxidation genes in several coral polyps. While we did not directly detect H2S in the GVC, active sulfur cycling may still occur within the GVC microenvironment, e.g. due to close spatial association between sulfate reducing and sulfide oxidixing bacteria and/or efficient chemical sulfide oxidation in the GVC. Together, these observations indicate that the GVC functions as an important site of microbial activity, which may play a major role in the internal nutrient cycling of the coral holobiont.

Our H2 measurements after feeding reflect dynamic changes in the availability of organic matter within the GVC. In *C. curvata* polyps, our observation showed that H2 concentrations declined to undetectable levels after 36–48 hours, indicating low levels of residual organic substrates. In *Goniastrea* polyps, measurements started immediately after feeding, and H2 concentrations rose rapidly within 10 minutes. This sharp increase suggests a rapid accumulation of labile organic carbon in the GVC following feeding, driven by coral digestive processes and microbial decomposition. As microbial carbon metabolism is triggered by the availability of labile organic carbon (Görke and Stülke, 2008), we speculate that nitrogen cycling is similarly influenced by nitrogen-containing organic compounds and exhibits feeding-induced dynamics, as observed e.g. in the vertebrate gut (Heinken et al., 2012; Rowland et al., 2017; Herrera and Byerley, 2025). In this study, we performed a single artificial feeding. However, under natural conditions, microbial activity in the GVC likely fluctuates with feeding frequency, food composition, digestive efficiency, and the retention time of food residues, leading to strong temporal variability in both carbon turnover and nitrogen transformations.

When comparing the GVC microbial communities of polyps with consistently low NO concentrations (i.e., Healthy Polyp 2 and Bleached Polyp 1) with other polyps, we found that no taxa with predicted denitrification genes were detected in the low-NO group, while taxa with denitrification potential were detected in every high-NO sample. Additionally, several bacterial families showed clear shifts in relative abundance. High-NO polyps were enriched in Xanthobacteraceae, a family which includes many denitrifiers and has been reported to perform partial denitrification (Gómez et al., 2005; Zhou et al., 2023), and Alcanivoracaceae, which also possess denitrification potential (Chen et al., 2024). In the low-NO polyps, Pseudoalteromonadaceae were more abundant. This family is characterized by strong protease production and organic nitrogen degradation (Chen et al., 2020), and only a few species within this family are capable of denitrification (Ivanova et al., 1998). We found that the relative abundance of predicted *nifH* genes in our coral polyps was much lower than similar predictions applied to other coral microbiomes (e.g. Lesser et al. 2017). However, these predictions do not reflect the true abundance of functional genes, and since we detected H2 production under low O2 conditions we cannot rule out a potential contribution of diazotrophs to the H2 dynamics. In addition to species-specific differences, this difference may also reflect differences in nitrogen sources or availability in the surrounding environment. Therefore, variation in microbial community composition may be accompanied by shifts in nitrogen transformation pathways within the GVC environment.

In bleached corals, the loss of symbiotic algae reduces the primary energy supply for the host, which consequently has to rely more on heterotrophic feeding and changes in the composition of ingested food (Grottoli et al., 2006; Houlbrèque and Ferrier-Pagès, 2009; Meunier et al., 2019). Increased coral heterotrophic feeding may enhance microbial activity in the GVC. However, in this study, we did not quantitatively control the amount of food provided or the coral polyp size, and therefore cannot directly compare substrate concentrations to infer different reaction rates. In addition, NO and N2O are intermediate products of denitrification and do not fully represent the overall process; applying the acetylene inhibition technique, which blocks the conversion of N2O to N2, before N2O measurements would provide a more accurate quantification of denitrification activity (Yoshinari et al., 1977; Sørensen, 1978). Finally, our study only assessed taxonomic composition and taxonomy-based functional prediction in the GVC microbiome. While these predictions can contribute to meaningful hypothesis generation, a metagenomic or metatranscriptomic approach is required for true functional profiling. Future studies that incorporate controlled feeding experiments, measurements of carbon and nitrogen contents in the GVC, in concert with metabolic gene profiling will be essential for resolving the role of the GVC-associated microbiome in holobiont C–N cycling.

In summary, this study demonstrates that the coral gastrovascular cavity is a chemically dynamic microenvironment and a hotspot of microbial activity. Based on microsensor measurements of H2, NO and N2O, we show that the GVC supports active anaerobic microbial processes such as fermentation and denitrification, and we also identified corresponding microbial taxa and predicted functional genes within the GVC microbiome. In healthy corals, these anaerobic activities are suppressed under illumination by the hyperoxic conditions generated through symbiont photosynthesis. In contrast, the GVC of bleached corals remained consistently hypoxic and acidic, allowing anaerobic metabolic processes to persist across the diel cycle. Such patterns indicate a shift in energy and nutrient cycling pathways within the bleached coral holobiont. Furthermore, feeding experiments confirmed that the availability of organic substrates is an important driver of H2 dynamics, indicating that coral feeding behavior can strongly influence microbial activity within the GVC. Collectively, these findings establish the coral GVC as a critical site of microbial activity that is closely linked to host physiology and stress status. Future quantitative studies integrating microbial, physiological, and biogeochemical perspectives will be essential for resolving its contributions to internal nutrient cycling and to holobiont functioning under environmental change.

## Acknowledgements

This study was supported by a grant from the Gordon and Betty Moore Foundation (grant no. GBMF9206; https://doi.org/10.37807/GBMF9206) to MK, and a PhD scholarship from the Chinese Scholarship Council (CSC) to QZ. We acknowledge excellent technical assistance for coral husbandry by Mikkel Hansen and Sofie L. Jakobsen. The technical staff of Heron Island Research Station is thanked for excellent technical assistance and guidance.

