## Supplementary material for "Chemical Microenvironment of the Coral Gastrovascular Cavity and Its Associated Bacterial Diversity": Table S1, Figure S1 and Figure S2

**SUPPLEMENTARY INFORMATION**

**Table S1.** Microbial functions and corresponding KEGG Orthology (KO) IDs.

| Function | Enzyme | KEGG Orthology (KO) ID |
| --- | --- | --- |
| Antioxidant enzyme catalase | catalase | ko:K03781 |
| Denitrification |  | ko:K00370, ko:K00371, ko:K00374, ko:K02567, ko:K02568, ko:K00368, ko:K15864, ko:K04561, ko:K02305, ko:K00376 |
| Nitrogen fixation | nifH | ko:K02588 |
| H_2_ production | ferredoxin hydrogenase (EC 1.12.7.2), | ko:K00532, ko:K00533, ko:K00534, ko:K06441, ko:K18016, ko:K18017, ko:K18023 |
|  | iron hydrogenase (EC:1.12.7.-) | ko:K25123, ko:K25124 |
|  | bifurcating [FeFe] hydrogenase (EC 1.12.1.4) | ko:K17997, ko:K17998, ko:K17999 |
|  | [NiFe] hydrogenase | ko:K18008, ko:K00437 |
|  | formate hydrogenlyase | ko:K15827, ko:K15828, ko:K15829, ko:K15830, ko:K15831, ko:K15832, ko:K22015 |
|  | hydrogenase (EC 1.12.99.6) | ko:K06281, ko:K06282, ko:K23548, ko:K23549 |
|  | hydrogenase (EC 1.12.1.2) | ko:K00436, ko:K18005, ko:K18006, ko:K18007 |
|  | hydrogenase (EC 1.12.1.3) | ko:K17992, ko:K17993, ko:K17994, ko:K18330, ko:K18331, ko:K18332 |
|  | nitrogenase (EC 1.18.6.1) | ko:K00531, ko:K02586, ko:K02588, ko:K02591 |
|  | vanadium-dependent nitrogenase (1.18.6.2) | ko:K22896, ko:K22897, ko:K22898, ko:K22899 |
| H_2_ consumption | quinone-reactive Ni/Fe-hydrogenase (EC:1.12.5.1) | ko:K05922, ko:K05927 |
|  | acetogenesis | ko:K00192, ko:K00195, ko:K00196, ko:K00198 |
|  | sulfhydrogenase (EC:1.12.1.3, 1.12.1.5) | ko:K17993, ko:K17994, ko:K17995, ko:K17996 |
| Sulfate reduction |  | ko:K11180, ko:K11181, ko:K27196, ko:K27187, ko:K27188, ko:K27189, ko:K27190, ko:K27191 |
| Sulfide Oxidation |  | ko:K17222, ko:K17223, ko:K17224, ko:K17225, ko:K17226, ko:K17227, ko:K22622, ko:K17218, ko:K21307, ko:K21308, ko:K21309 |

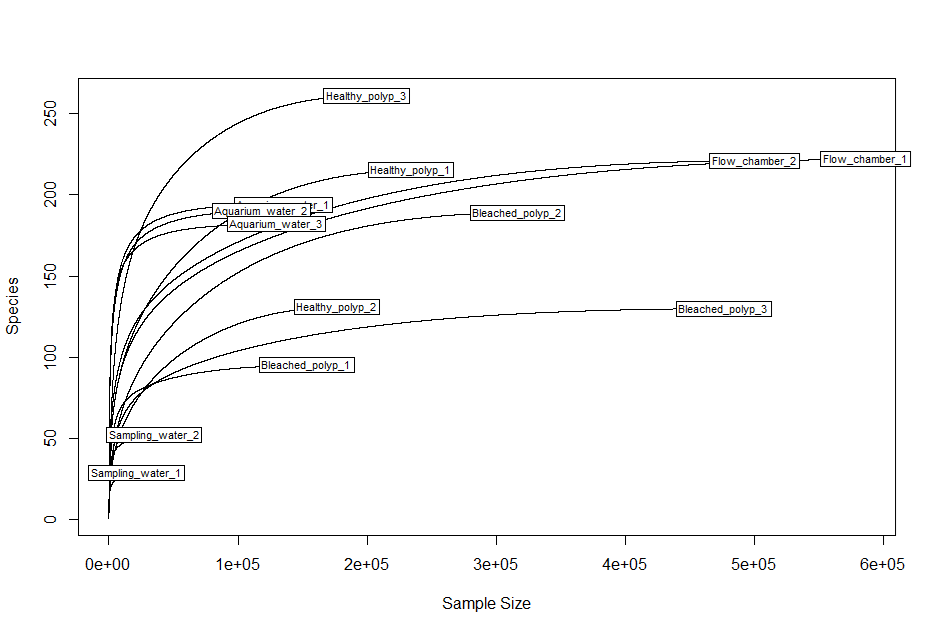

Figure S1 Rarefaction curves showing the relationship between sequencing depth and number of ASVs for each sample.

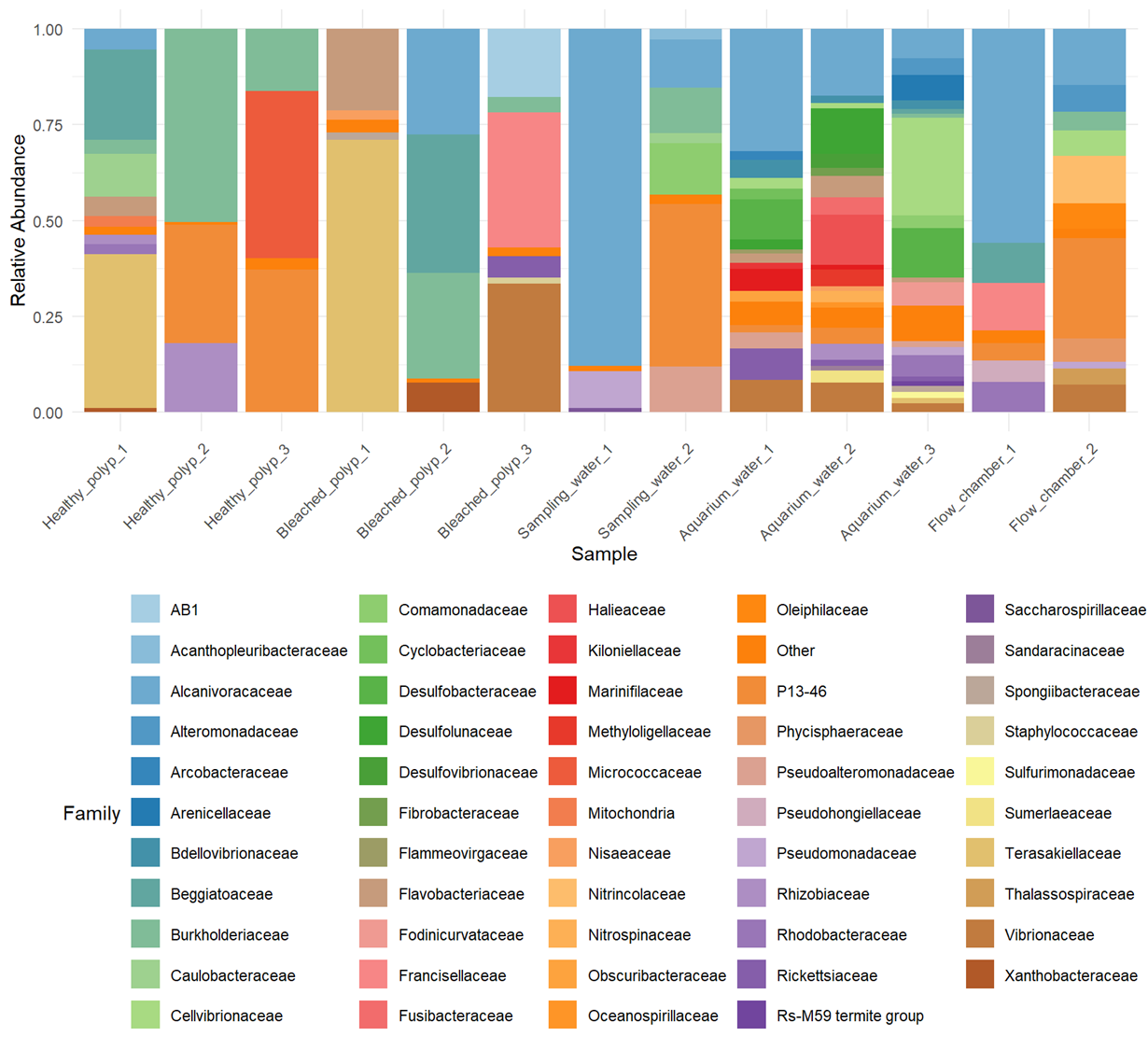

Figure S2 Relative abundance of bacterial families in the gastrovascular fluid of healthy and bleached *Caulastrea* *curvata* polyps and in environmental samples. Taxa with relative abundances below 1% across all samples were grouped as “Other”.
